# Unexpected distribution of the 4-formylaminooxyvinylglycine (FVG) biosynthetic pathway in *Pseudomonas* and beyond

**DOI:** 10.1101/2021.02.08.430221

**Authors:** Edward W. Davis, Rachel A. Okrent, Viola A. Manning, Kristin M. Trippe

## Abstract

The biological herbicide and antibiotic 4-formylaminooxyvinylglycine (FVG) was originally isolated from several rhizosphere-associated strains of *Pseudomonas fluorescens*. Biosynthesis of FVG is dependent on the *gvg* biosynthetic gene cluster in *P. fluorescens*. In this investigation, we used comparative genomics to identify strains with the genetic potential to produce FVG due to presence of a *gvg* gene cluster. These strains primarily belong to two groups of *Pseudomonas, P. fluorescens* and *P. syringae*, however, a few strains with the *gvg* cluster were found outside of *Pseudomonas*. Mass spectrometry confirmed that all tested strains of the *P. fluorescens* species group produced FVG. However, *P. syringae* strains did not produce FVG under standard conditions. Several lines of evidence regarding the transmission of the *gvg* cluster including a robust phylogenetic analysis suggest that it was introduced multiple times through horizontal gene transfer within the *Pseudomonas* lineage as well as in select lineages of *Thiomonas, Burkholderia* and *Pantoea*. Together, these data broaden our understanding of the evolution and diversity of FVG biosynthesis. In the course of this investigation, additional gene clusters containing only a subset of the genes required to produce FVG were identified in a broad range of bacteria, including many non-pseudomonads.

## INTRODUCTION

Members of the genus *Pseudomonas* are prolific producers of specialized metabolites that facilitate life in their environment. These chemicals contribute to many processes, including nutrient acquisition, host colonization, manipulation of host physiology and competition with other microbes. One example of a specialized metabolite produced by some strains of *Pseudomonas fluorescens* is 4-formylaminooxyvinylglycine (FVG, **Fig. 1a**), a non-canonical amino acid [1,2]. FVG arrests the germination of weedy grasses, and thus was originally termed a germination-arrest factor (GAF) [3]. FVG also displays activity against the bacterial plant pathogen *Erwinia amylovora*, the causal agent of fireblight [4]. As with other vinylglycines, the activity of FVG is attributed to inhibition of enzymes that utilize pyridoxal phosphate as a co-factor, such as 1-aminocyclopropane-1-carboxylate synthase in the ethylene biosynthesis pathway of plants and several enzymes involved in amino acid metabolism in bacteria [5,6]. The relatively recent discovery of FVG contrasts with the identification of other vinylglycines of bacterial origin that occurred decades earlier, e.g. rhizobitoxine [7,8], aminoethoxyvinylglycine (AVG) [9], and 4-methoxyvinylglycine (L-2-amino-4-methoxy-*trans*-3-butenoic acid) (MVG/AMB) [10].

**Fig. 1.**
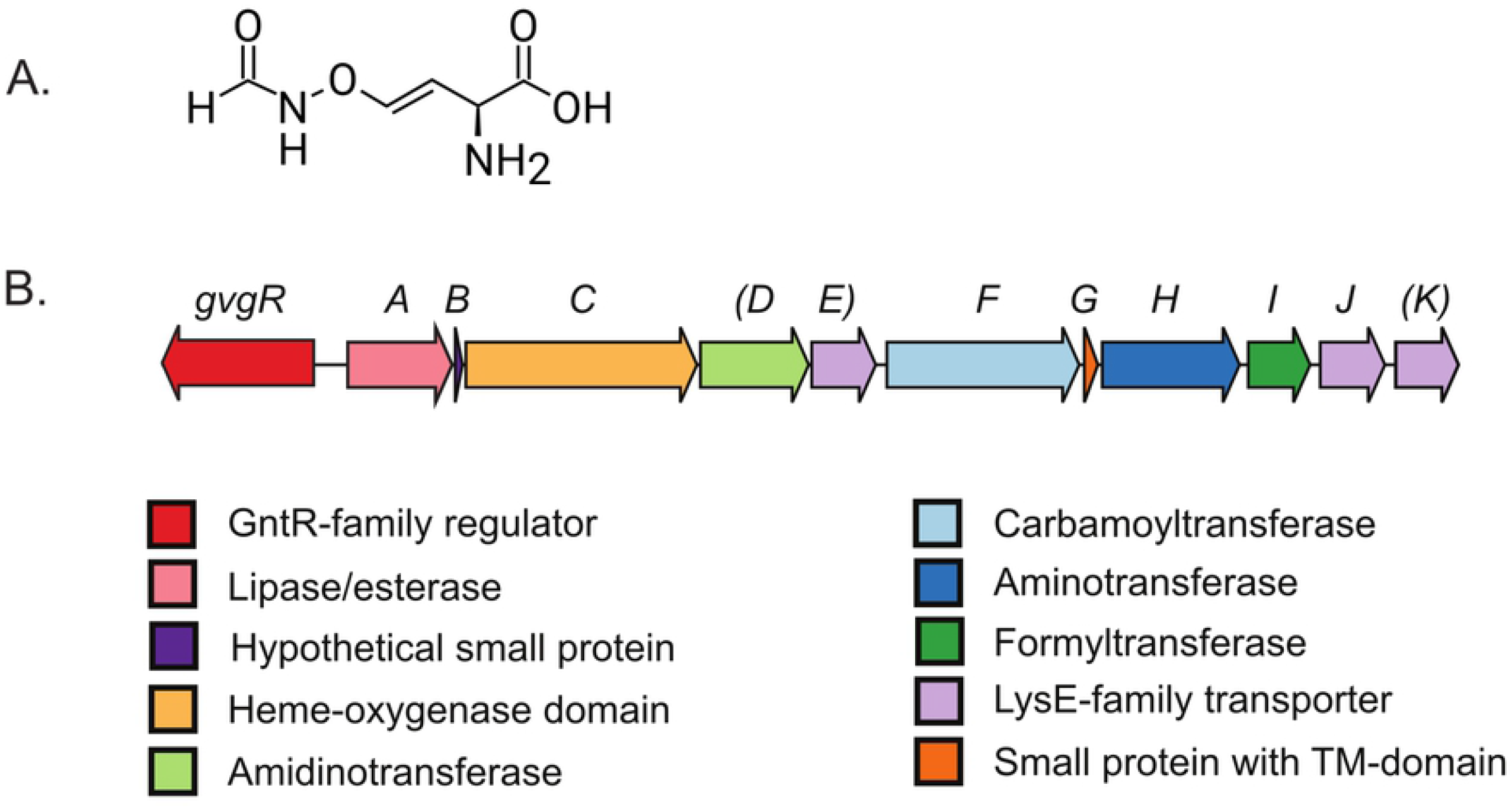
Structure of the 4-formylaminooxyvinylglycine compound and the gene cluster necessary for its production. **(a)** Structure of 4-formylaminooxyvinylglycine (FVG). **(b)** Organization of the *gvg* gene cluster from *Pseudomonas fluorescens* WH6. Arrows representing genes are colored based on the function of the encoded proteins. Gene notations in parentheses are not required for FVG production.

In earlier studies, FVG production was restricted to a small number of *P. fluorescens* isolated from grass and wheat rhizosphere in the Willamette Valley of Oregon [11]. FVG production was not detected in a phylogenetically diverse group of other *P. fluorescens* strains, including *P. fluorescens* A506, PfO-1, Pf-5, and SBW25 [2,3,12]. The narrow host range and geographic origin of FVG-producers coupled with the relatively recent identification of FVG might suggest that it is produced by few strains in a common, but specialized, ecological niche. However, our understanding of the relationship between these strains is incomplete. Previous analyses included a limited number of strains, either due to lack of available genomic or gene sequences [13], or lack of available cultures for bioassays [3]. As the available genome sequences in the *Pseudomonas fluorescens* group has greatly expanded, along with the sequences of the genus as a whole, a more robust analysis using phylogenomic and targeted locus scrutinization is now possible [14].

The 13-kb GAF vinylglycine (*gvg)* biosynthetic gene cluster (**Fig. 1b)** is essential for FVG production in *P. fluorescens* WH6 [15–17], and the presence of this cluster is correlated with the ability of other *P. fluorescens* strains to produce FVG [2]. Although the biosynthetic pathway for FVG has not yet been elucidated, the genes within the *gvg* cluster required for FVG production have been determined in *Pfl* WH6 [17]. The *gvg* cluster has also been reported in genomes of a few other species. For example, several genes similar to those of the *gvg* cluster were sequenced and characterized in *P. chlororaphis (Pch)* PCL1391, though in relation to the regulation of phenazine rather than FVG [18]. *P. syringae* (*Psy*) pv. *maculicola* ES4326, also known as *P. cannabina (Pca)* pv. *alisalensis* ES4326, was reported to contain the *gvg* cluster but not to produce FVG under standard conditions [16]. Screens for novel antibiotics recently revealed that the non-pseudomonad *Pantoea ananatis (Pan)* BRT175 encodes the *gvg* biosynthetic gene cluster [19]. These examples suggest a more complex history of FVG biosynthesis but is not a systematic analysis.

In this study, we combine the analysis of the original collection of Willamette Valley FVG-producers with mining of publicly available sequence databases to define the phylogenetic, host and geographic distribution of the *gvg* cluster. We also determine if the presence of the *gvg* cluster is sufficient for FVG production in distinct genetic backgrounds. The origin of the *gvg* cluster in various strains, through vertical descent or horizontal gene transfer, is explored.

We use a robust phylogenetic analysis to test the hypothesis that the *gvg* locus is restricted to a monophyletic clade of *Pseudomonas* spp. In the course of the analysis, additional “orphan” biosynthetic gene clusters are identified that share similarities with the *gvg* cluster but are predicted to produce distinct compounds.

## METHODS

### Search for strains with *gvg* clusters

The formyltransferase gene was used as a probe in two general strategies to search for sequenced strains containing the *gvg* biosynthetic gene cluster. The NCBI database was searched for homologs of the formyltransferases encoded by *gvgI* from *Pfl* WH6 (PFWH6_5257) and *Pan* BRT175 (L585_04235) using BLAST+ 2.3.0 (BLASTP and TBLASTN) under default parameters [20]. Locus tags were retrieved from positive hits and used to search the NCBI Nucleotide database to confirm the presence of the other genes in the cluster. The Gene Neighborhood tool in Integrated Microbial Genome (IMG) v4 from the Joint Genome Institute was also used to search for complete gene clusters, again with the *gvgI* as a probe. Comparisons of the gene content or genetic context were made using the Gene Neighborhood tool or manually based on gene annotations. Sequences were downloaded from NCBI or IMG and imported into CLC Main Workbench v. 6 (Qiagen) for additional analysis. For synteny comparisons, chromosome regions were cropped approximately 25 kb upstream and downstream of the CTase-encoding gene and colored based on the synteny of genetic contexts.

Variations of the *gvg* cluster that may lack homologs of *gvgI*, termed *gvg*-like clusters, were identified using the *gvgA* (PFWH6_5249), *gvgC* (PFWH6_5251), and *gvgF* (PFWH6_5254) genes. The NCBI database was searched for homologs of the amino acid sequences corresponding to these genes from *Pfl* WH6 by BLASTP. Additional homologs were retrieved by Genome BLASTP in IMG. The Gene Neighborhood tool in IMG was used to facilitate identification of gene clusters.

### Phylogenetic tree construction of the *P. fluoresens and P. syringae* groups

The multi-locus sequence analysis based phylogenetic tree was generated as previously described [21]. The code for the automated multi-locus sequence analysis pipeline (automlsa2) is available from the python package index (pypi) at https://pypi.org/project/automlsa2/ and builds upon autoMLSA.pl as previously reported [22]. Briefly, TBLASTN+ (v. 2.10.0) was used to identify and extract sequences for 99 of the 100 housekeeping genes as identified in Hesse et al. 2018 [14]; PA4393 was excluded from our analysis as it was unable to be routinely recovered using TBLASTN+ from all selected Pseudomonas spp. Included genome sequences are shown in **Table S1**. As *P. aeruginosa* has previously been shown to be a suitable outgroup for the *P. fluorescens* and *P. syringae* groups, three selected *P. aeruginosa* isolates were used as an outgroup [23]. All genes that passed filter were aligned using MAFFT (v. 7.427) [24,25].

Phylogenetic tree construction was done using IQ-TREE2 [26–29] with a partitioned nexus file containing references to each aligned sequence as input. Model selection and partition finding was done using the ‘-m MFP --merge rclusterf’ mode in IQ-TREE2, with protein models limited with the ‘--msub nuclear’ flag. Branch supports were determined using 1000 ultra fast bootstraps, and 1000 SH-aLRT test replicates (-B 1000, -alrt 1000). Nodes with >=95% and >=80% supports, respectively, were considered as well supported, and are so marked on the trees. Nodes with less support are unlabeled. Trees were visualized using ggtree v2.4.1 in R v4.0.1[30–33].

In addition, the *P. fluorescens* and *P. gessardii* subgroups were trimmed from the larger *Pseudomonas* phylogeny and annotated using ggtree, as above, and inkscape (https://inkscape.org/). The genes present at a particular locus, between the genes encoding a methyl transferase and single-stranded DNA binding protein (*tam* and *ssb)*, were identified for these strains. Identification was performed by searching for the sequence encoded by the *P. fluorescens tam* gene using IMG BLASTP (v4), cross-referencing results with *P. fluorescens* and *P. gessardii* subgroups in the phylogeny, and manually examining gene annotations. A phylogenetic tree including *Pseudomonas* strains encoding the gvg-gene cluster was constructed generally as above, but with *Pan* BRT175 as the outgroup.

### Gene diagram for *gvg* locus

The *gvg* locus in each genome was identified using BLAST with the protein sequences from WH6 used as queries against the predicted protein sequences from each of the target genomes as the databases. Subsetted genbank files (accessions listed in the figure) with 10kb regions upstream and downstream the putative *gvg* loci were extracted for each representative genome (20kb in total). Subset genbank files were used as input to clinker for visualization [34].

### Comparison of tree topologies

One strain from each clade in the *Pseudomonas* phylogeny containing the *gvg* cluster was selected for comparison of the topologies of *gvg* and *species* trees with *Pan* BRT175 as the outgroup. Seven of the Gvg proteins (GvgR, GvgA, GvgC, GvgF, GvgI, and GvgJ) were used in construction of the *gvg* phylogeny. Tree topology tests, including the approximately unbiased (-au) and Shimodaira-Hasegawa (SH) tests were done using iqtree2 v2.1.2 with 10,000 replicates (-zb 10000) [35,36].

### Phylogenetic tree of carbamoyltransferase proteins

Carbamoyltransferase (CTase) sequences were retrieved using BLASTP (v2.3.0) of the NCBI nt database with PFWH6_5254 as the query sequence. Additional sequences were identified by a genome BLASTP search of the IMG database (v4.540). Sequences were confirmed to be present within clusters containing other homologs to *gvg* cluster genes. CTase sequences from the in-house genomes were added and all sequences were aligned using MAFFT (v. 7.244 [24]) and the percent identities calculated in CLC Main Workbench. A representative sequence of groups of sequences with greater than 95% identity was chosen for further analysis. These sequences were aligned again using MAFFT with the L-INS-I setting and trimmed using Gblocks (v. 0.91b) with half gapped positions allowed (-b5 = h setting). The LG substitution model was selected based on the PROTGAMMAAUTO option in RAxML (v8 [37]). Bootstrap analysis was performed using the autoMRE bootstrapping criterion, resulting in 350 bootstrap replicates. The resulting phylogenetic tree was visualized and annotated in iTOL [38].

### Bacterial isolation

Bacterial strains used in this study are listed in **Table 1**. The strains in the ARS collection of FVG producers were isolated from wheat and grass rhizosphere in the central Willamette Valley of OR, USA as described in [11]. *Pseudomonas fluorescens* P5A was isolated from the surface of a basidiomycete fungus, *Marasmius oreades*, at the edge of a Douglas fir plantation in the Willamette Valley. The sample was transferred to sterile water (100 ml) with glass beads and shaken. The resulting suspension was serially diluted, plated on Fluorescent Pseudomonad Media (FPM) agar with cycloheximide (75 mg/L), and incubated at 28 °C for 24 h. Colonies were re-streaked on FPM agar and screened for germination arrest and anti-*Erwinia* activity consistent with FVG production as in [3,4]. Initial taxonomic identification of *P. fluorescens* P5A was performed by amplifying and sequencing 16S rDNA using standard protocols.

**Table 1.** Strains used in this study.

| Strain* | Origin or description | Location | Reference | Genome Reference |
| --- | --- | --- | --- | --- |
| *Pch* 30-84 | Wheat rhizosphere | WA, USA | [61] | [46] |
| *Pch* O6 | Soil | UT, USA | [62] | [46] |
| *Pfl* A3422A | Unknown | OR, USA | [2] | [39] |
| *Pfl* AH4 | Bluegrass rhizosphere | OR, USA | [3] | [39] |
| *Pfl* E24 | Bluegrass rhizosphere | OR, USA | [2] | [39] |
| *Pfl* G2Y | Ryegrass rhizosphere | OR, USA | [2] | [39] |
| *Pfl* P5A | Fungal surface | OR, USA | This study | [39] |
| *Pfl* WH6 | Wheat rhizosphere | OR, USA | [11] | [15] |
| *Pfl* WH6-30G | $\Delta gvgH$ mutant | N/A | [17] | N/A |
| *Pfl* WH6-31G | $\Delta gvgI$ mutant | N/A | [17] | N/A |
| *Pfl* TDH40 | Bluegrass rhizosphere | OR, USA | [13] | [39] |
| *Psn* A342 | Unknown | OR, USA | [13] | [39] |
| *Psn* BG33R | Peach rhizosphere | SC, USA | [63] | [46] |
| *Pca* BS91 | Broccoli raab | CA, USA | [64] | [65] |
| *Psy* ES4326 | Radish | WI, USA | [66] | [67] |
| *Pan* BRT175 | Strawberry | Unknown | [19] | [68] |
| *Eam* 153 | Apple | OR, USA | [69] | N/A |

\**Eam*, *Erwinia amylovora*; *Pan*, *Pantoea ananatis*; *Pca*, *Pseudomonas* pv. *alisalensis*; *Pch*, *Pseudomonas chlororaphis*; *Pfl*, *Pseudomonas fluorescens*; *Psp*, *Pseudomonas* sp.; *Psn*, *Pseudomonas synxantha*; *Psy*, *Pseudomonas syringae*.

The sequencing, assembly and annotation of the genomes is described in Okrent et al. [39]. The contigs were ordered based on whole genome alignment with closely-related complete genomes and the presence of the *gvg* gene cluster was confirmed.

### Preparation of culture filtrates and biological assays

Bacteria were inoculated into either 60 ml or 5 ml of Pseudomonas Minimal Salts Medium [3] in a 125-ml Wheaton bottle or 20-ml test tube, respectively. The cultures were grown at 28 °C with shaking for either seven (60-ml cultures) or two days (5-ml cultures). Subsequently, cells were removed from the filtrate by centrifugation (3000 x g for 15 min) and filter sterilization (Millipore GP express Steritop, 0.22-µm pore size (EMD Millipore) or 25-mm Pall Acrodisc with 0.22-µm pore size Supor membrane (Pall Corporation), respectively. For initial LAESI-MS experiments, culture filtrate was extracted with90% ethanol as described previously [1].

The agar diffusion bioassays for anti-microbial activity against *Erwinia amylovora* were performed generally as described in [4], however smaller volumes were used than reported previously. Aliquots of *E. amylovora* cultures grown overnight and diluted to an OD600 of 0.2 (300 µL) were spread on a 100×100×15 mm square plate containing 35 mL of 925 agar media. Four wells per plate were cut using a #2 cork-borer and the plugs removed. Samples of filtrate (40 µL) were added per well and plates incubated at 28 °C for 2 days. Plates were scanned and the areas of the zones of inhibition were measured on triplicate wells for each sample using ImageJ from the National Institutes of Health [40].

### LAESI-MS analysis

LAESI-MS analysis was performed by Protea Biosciences, Inc. Samples of crude culture filtrate, non-inoculated filtrate, or extracted culture filtrate (20 µl) were aliquoted into individual wells of a 96-well plate and analyzed with a LAESI DP-1000 (Protea Biosciences) coupled to a Q Exactive Orbitrap mass spectrometer (Thermo Scientific) according to the methods described in [41].

Conditions were slightly altered for analysis of FVG in crude culture filtrates. Settings for the LAESI DP-1000 were electrospray (+ 4000 V, 1.0 µl/min flow, 50% methanol, 0.1% acetic acid), 100 pulses per well at 20 Hz laser repetition rate, and laser energy ∼600 µJ. The samples were analyzed in selected-ion monitoring mode at a resolution of 140,000 with scans centered on the sodium adduct of FVG, [M+Na]^+^ *m/z* 183.037. The average scan signal intensities were extracted on each well using LAESI Bridge software. Measurements were also performed on a second set of biological replicates for comparison.

## RESULTS AND DISCUSSION

### Identification of potential FVG-producers

The *gvg* gene cluster in *Pfl* WH6 contains 12 genes (*gvgR, gvgA*-*gvgK*), encoding putative regulatory, transferase or transport activity, and two small ORFs encoding peptides of less than 50 amino acids **(Fig. 1b)** [16,17]. Three genes, *gvgD, gvgE* and *gvgK*, are dispensable for production of FVG [17]. In preliminary analyses, the *gvgI* gene, encoding a formyltransferase, was proposed as a specific marker for FVG biosynthesis [13]. Based on previous studies, we hypothesized that the *gvgI* gene would be found in a limited number of isolates within the three previously reported groups of *Pseudomonas* (i.e. WH6-like strains, *P. chlororaphis*, and few *P. syringae* isolates), in addition to *Pantoea ananatis* isolate known to encode the *gvg* locus.

Our strategy for prediction of additional FVG-producing strains was to mine genome databases for the *gvgI* gene and confirm its presence within a *gvg* cluster. This strategy identified over 30 additional sequenced strains containing a *gvg* cluster in NCBI and IMG databases as of December 2020. Counter to our expected result, the strains originated from a broad spectrum of geographic locations and varied taxonomic units **(Table S2)**. In general, strains encoding the *gvg* cluster were collected from rhizosphere or soil samples and belong to the genus *Pseudomonas*. However, the *gvg* cluster was also found in other genera including *Thiomonas, Pantoea* and *Burkholderia*, and strains from environmental or clinical samples **(Table S2)**.

### Distribution of *gvg*-cluster containing strains

The distribution of strains containing the *gvg* cluster within the genus *Pseudomonas* is not readily apparent by species name alone. The *Pseudomonas* genus contains over 230 recognized species names (**www.bacterio.net/pseudomonas.html)**. The *P. fluorescens* and *P. syringae* groups or species complexes are diverse and heterogeneous, and are each further divided into subgroups. Recent phylogenies have grouped *P. fluorescens* into 9 subgroups [42,43] and *P. syringae* into at least 9 genomospecies or 11 phylogroups with multiple subgroups [44]. As a further complication, several of the *Pseudomonas* strains containing a *gvg* cluster have not been conclusively assigned to species **(Table S2)**. Additionally, researchers have suggested that some species designations in *Pseudomonas* may be incorrect based on newer tools, namely comparisons to type strains using multilocus or whole genome-based methods [14,42,45].

As the species labels on these isolates do not allow us to make conclusions about phylogenetic relationships between them, and in order to investigate the distribution of strains containing the *gvg* cluster within the genus *Pseudomonas*, a maximum likelihood phylogenetic tree was constructed from the amino acid sequences of 99 housekeeping genes [14,46], focusing on groups containing strains with *gvg*-like clusters. The resulting phylogenetic tree, shown in **Fig. 2A**, is largely consistent with the phylogenetic tree published in Hesse et al. 2018 [14,42,43,47] and clustering of average amino acid identities from whole genomes [14,48]. Strains encoding the *gvg* cluster are shown using blue or green circles at the tips for the *P. fluorescens* and *P. syringae* groups, respectively.

**Fig. 2.**
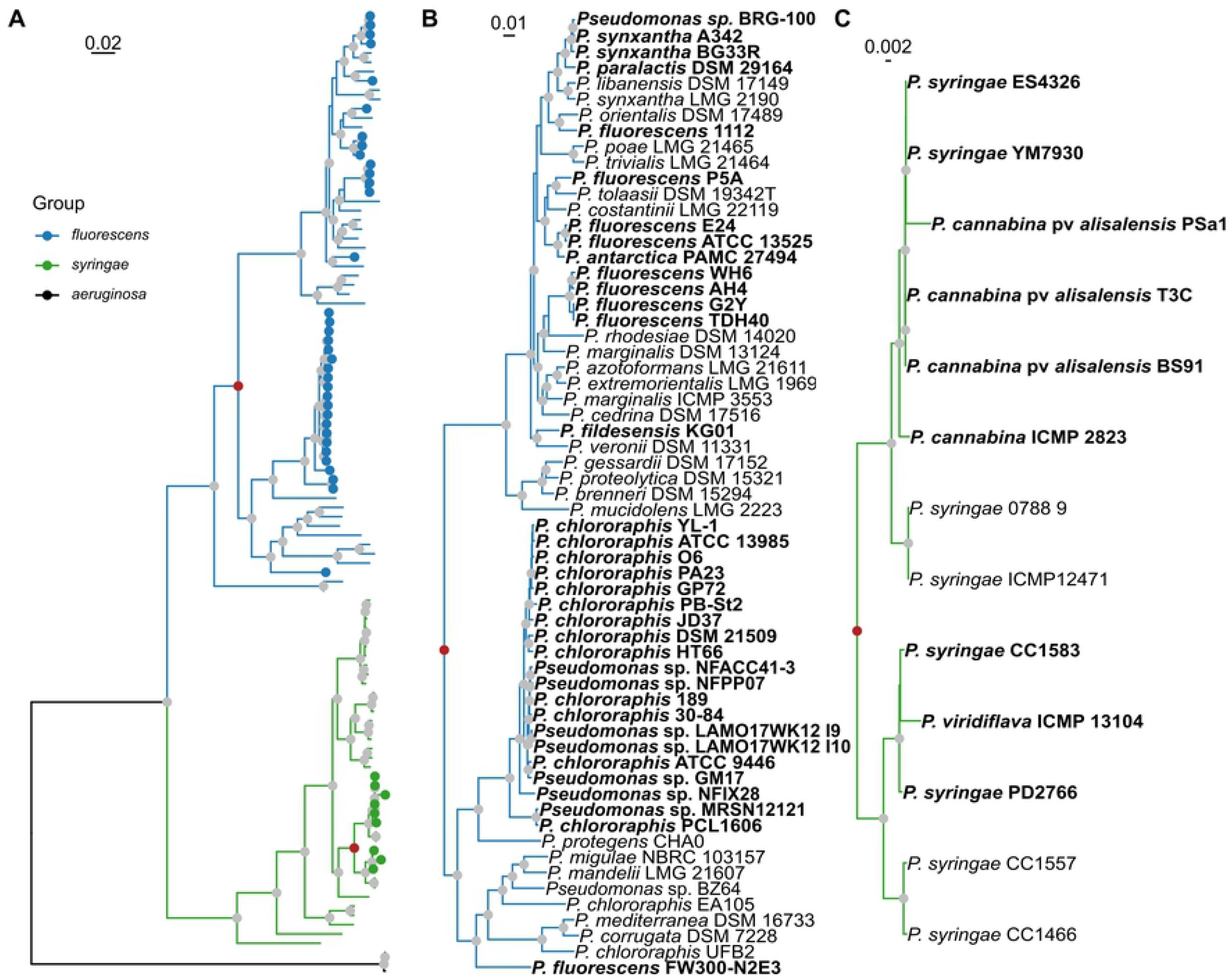
The *gvg* locus has a patchy distribution in genomes within the *Pseudomonas* genus. (A) Full maximum likelihood phylogenetic tree based on 99 housekeeping genes. The *fluorescens* and *syringae* groups are highlighted in blue and green, respectively. Red nodes correspond to the expanded clades in B and C. (B) Expanded *P. fluorescens* clade. (C) Expanded *P. syringae* clade. The tree was rooted on *P. aeruginosa*. Strain names in bold contain a *gvg* cluster. Nodes indicated with a gray circle are well supported (>95% UF-bootstraps & >80% SH-aLRT replicates). Genome accessions are listed in **Table S1**.

Despite the known limitations of accurate taxonomic designations within the *Pseudomonas* genus, analysis of the *Pseudomonas* phylogeny revealed that strains with homologous *gvg* clusters are present in the *P. fluorescens* and *P. syringae* groups only. Further, and contrary to our original hypothesis, the gene clusters appear in multiple subgroups within the *P. fluorescens* group, i.e. the *P. fluorescens* and *P. chlororaphis* subgroups, as well as *Pfl* FW300-N2E3, not yet placed in a subgroup. Strains containing the cluster within the *P. fluorescens* subgroup are not monophyletic, but instead are dispersed throughout the subgroup **(Fig. 2B; bold)**. The strains from the original Willamette Valley collection of FVG producers are found in multiple clades within the *P. fluorescens* subgroup (**Fig. 2B; bold**). In contrast, *gvg* clusters are restricted to two paraphyletic groups in a small subclade of *P. syringae* **(Fig. 2C; bold)**. These strains are variously annotated as *P. cannabina* and *P. syringae* and include the model Arabidopsis pathogen *Psy* ES4326. The full phylogenetic tree is included as **supplemental data S1**.

The *gvg* cluster was also found in several non-pseudomonads, including one strain of *Pantoea ananatis, Thiomonas sp*. and several strains of *Burkholderia*. In each case, the presence of a *gvg*-like cluster is not typical of the species. For example, a PCR-based survey of 117 strains of *Pantoea* failed to detect the cluster in any strain other than *Pan* BRT175 [19]. Similarly, of the 22 sequenced strains of *Pantoea ananatis* in the IMG database as of December 2020, *Pan* BRT175 and the closely related *Pan* PANS 99-36 are the only strains with the cluster. The *gvg* cluster is also unique to *Thiomonas* sp. (*Tsp*) CB2 of the 11 strains of *Thiomonas* that have been sequenced. Likewise, although there were approximately 40 sequenced genomes in the *Burkholderia cepacia* complex, only three contain complete *gvg* clusters (**Table S2**).

### Gene content of *gvg* clusters

As *gvg* gene clusters are present in a variety of bacterial strains, we were interested in determining whether there are variations in the gene content. To this end, the clusters were examined for gene loss, gain or rearrangement. Rearrangement was not evident in these clusters, as genes consistently appear in the same order. However, gene loss and gain were both evident and consistent with the phylogeny, as shown in **Fig. 3**. The three genes not essential for FVG production, *gvgD, gvgE* and *gvgK*, are absent in some clusters. While most strains within the *P. fluorescens* subgroup have the same set of genes as *Pfl* WH6, the two strains in the E24 clade, *Pfl* E24 and *Pfl* ATCC 13525, as well as *Pfl* FW300-N2E3, lack the partially-redundant LysE transporter gene, *gvgK*. The gene cluster in *Pfl* FW300-N2E3 lacks segments of the non-essential amidinotransferase encoded by *gvgD*, indicative of gene decay. The strains within the *P. syringae* consistently lack *gvgK* but contain all other genes **(Fig. 3)**.

**Fig. 3.**
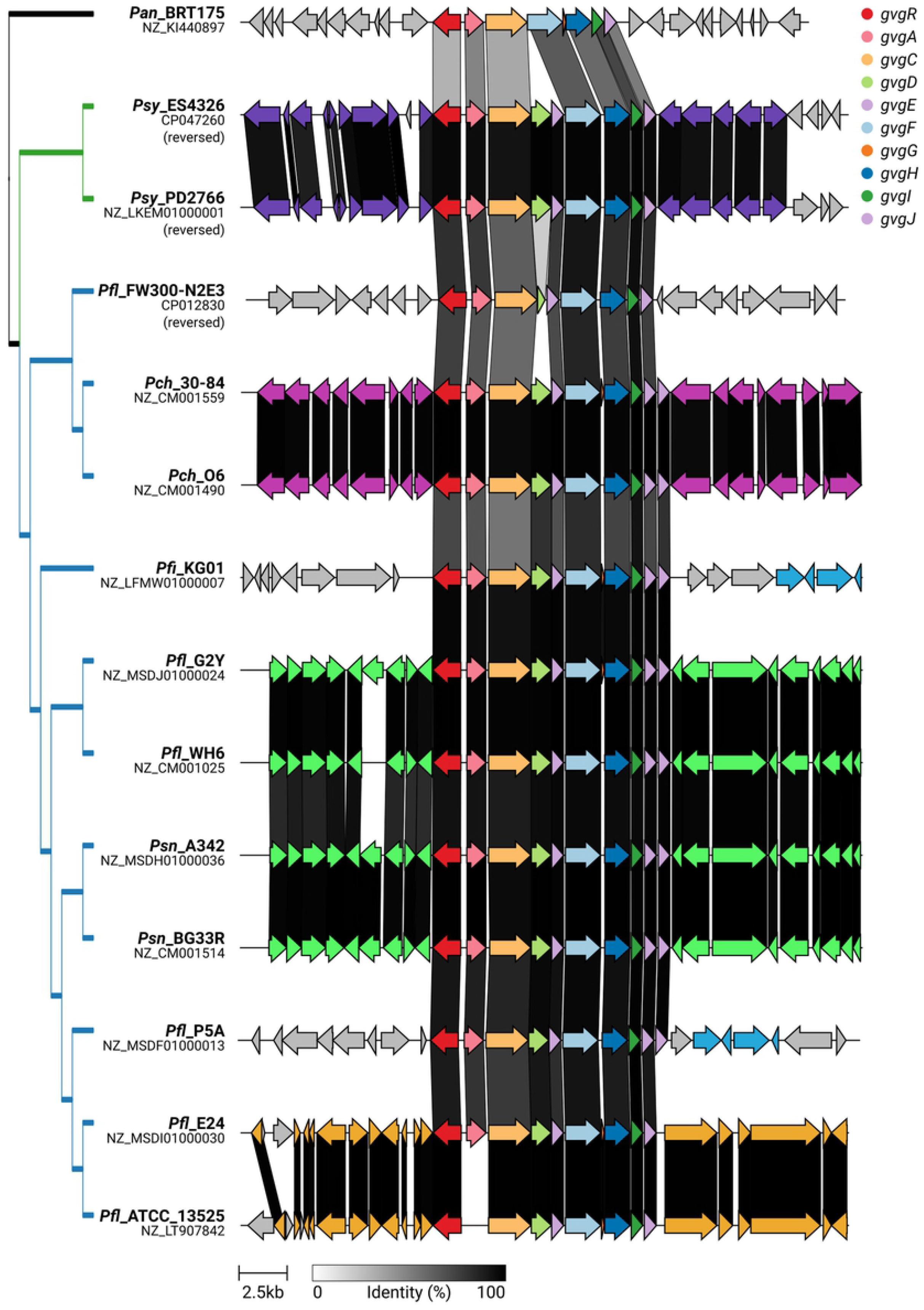
The *gvg* biosynthetic locus exhibits co-linearity but also gene gain and/or loss within *Pseudomonas* spp. The *gvg* clusters are color coded according to legend. Syntenic sets of neighboring genes are additionally shown in the same color (not listed in legend). Sequences shared between strains are connected by bands that are shaded according to percent identity (black = 100% identity, white = 0% identity). Strains are ordered by dendrogram on the left, which follows the species tree.

Outside of *Pseudomonas*, the *gvg*-like clusters are more variable. *Pan* BRT175 lacks orthologs of *gvgD, gvgE* and *gvgK* **(Fig. 3** and **Fig. S1**). *Tsp* CB2 lacks *gvgK* only and *Burkholderia* strains lack both extra LysE genes, *gvgD* and *gvgK* **(Fig. S1**). Uniquely, *Thiomonas* and *Burkholderia* clusters have gene sequences inserted between the *gvgR* and *gvgA* orthologs and between the *gvgH* and *gvgI* orthologs **(Fig. S1**). These genes encode putative amino acid transporters, a metallohydrolase, and proteins of unknown function. What effect the presence of these genes may have on the metabolites produced by these strains is unknown; they may produce a variant of FVG rather than the compound itself.

### Formylaminooxyvinylglycine chemical analysis

The routine determination of an FVG phenotype typically requires an ethanol-based extraction of crude culture filtrate followed by thin-layer chromatography [1] as well as two biological assays [3,4]. However, this protocol is not amenable to high-throughput analysis and is sometimes ambiguous. We recently developed a LAESI-MS-based method to directly detect FVG by mass in crude culture filtrates without extraction[41]. This protocol was used here to analyze a diverse selection of strains to determine if the presence of a *gvg*-like cluster corresponds to the ability to produce FVG. The culture filtrates were analyzed by LAESI-MS and, using *Erwinia amylovora* as an indicator strain, inhibition using the agar diffusion assay. All tested strains within the *P. fluorescens* group produced FVG, including strains in the *P. fluorescens* and *P. chlororaphis* subgroups **(Table 2)**. As previously reported, the non-pseudomonad *Pan* BRT175 also produced FVG **(Table 2)** [41]. However, neither biological activity consistent with FVG production nor the mass associated with FVG were detected from the three strains within the *P. syringae* group **(Table 2)**. The absence of FVG production from these strains is consistent with previous analyses of *Psy* ES4326 [16,19,41]. We investigated multiple strains from the *P. syringae* group in the current study, due to the possibility that a mutation in the laboratory strain *Psy* ES4326 was responsible for the lack of FVG production previously observed. This was not the case, as the closely related isolates of *Pca* also did not produce FVG.

**Table 2.** Zone of inhibition of bacterial filtrates against *Erwinia amylovora* and detection of FVG (m/z 183.037) by LAESI-MS for various strains.

| Strain* | Zone of inhibition<br>cm<sup>2</sup> (SD) | FVG<br>(M+Na)<br>183.037‡ |
| --- | --- | --- |
| *Pfl* WH6† | 6.79 (0.35) | + |
| *Pfl* AH4† | 6.60 (0.29) | + |
| *Pfl* TDH40 | 6.56 (0.27) | + |
| *Pfl* G2Y | 7.20 (0.32) | + |
| *Pfl* A3422A† | 6.36 (0.29) | + |
| *Pfl* E24† | 4.68 (0.18) | + |
| *Pfl* P5A | 4.82 (0.19) | + |
| *Psn* A342† | 5.11 (0.25) | + |
| *Psn* BG33R | 6.99 (0.24) | + |
| *Pch* O6 | 5.45 (0.31) | + |
| *Pch* 30-84 | 6.68 (0.22) | + |
| *Psy* ES4326† | 0.00 (0.00) | - |
| *Pca* BS91 | 0.00 (0.00) | - |
| *Pca* T3C | 0.00 (0.00) | - |
| *Pan* BRT175 | 7.78 (0.20) | + |

\**Pan*, *Pantoea ananatis*; *Pca*, *Pseudomonas* pv. *alisalensis*; *Pch*, *Pseudomonas chlororaphis*; *Pfl*, *Pseudomonas fluorescens*; *Psn*, *Pseudomonas synxantha*; *Psy*, *Pseudomonas syringae*.
† FVG production has previously been analyzed by different methods for these strains.
‡ Values are listed in **Table S3**.

Although we tested several possible hypotheses for the lack of FVG production in *P. syringae* strains, we were not able to conclusively demonstrate why these strains do not produce FVG under standard conditions. However, the most likely explanation is related to differences in gene regulation. *P. syringae* lack the sigma factor/anti-sigma factor pair involved in regulation of FVG production in WH6 encoded by the *prtIR* locus [49]. This suggests that regulation of FVG production differs in these strains relative to *P. fluorescens*. In *P. fluorescens* WH6, transcription of the *gvg* cluster occurs from multiple promoters, with one located in the intergenic region between *gvgR* and *gvgA* as well as additional promoters internal to the cluster [17]. We thus examined the promoter regions in *P. syringae* and compared them to those of several strains in the *P. fluorescens* group. The predicted promoter sequences located between *gvgR* and *gvgA* are conserved within *P. fluorescens* group strains (including *P. chlororaphis*) but substantially different in *P. syringae* strains **(Fig. S2)**. Therefore, *P. syringae* may require different conditions for induction of the *gvg* cluster, which may reflect the different ecological niches or nutrient acquisition strategies of the two species.

### Evidence of horizontal gene transfer

The observed pattern of *gvg* clusters, present in a given phylogenetic group but not closely related strains, is consistent with horizontal gene transfer (HGT) and/or the inclusion of clusters in genomic islands. Horizontally transferred genes typically demonstrate a lack of correspondence between phylogenetic trees representing gene evolution and those representing species evolution [50]. We therefore compared the phylogeny of the *gvg* genes and the housekeeping genes for strains within *Pseudomonas*. One representative from each clade that contains a *gvg* cluster was chosen for construction of maximum likelihood phylogenetic trees for amino acid sequences encoded by *gvg* and housekeeping genes. A comparison of the species tree and a partitioned Gvg protein maximum likelihood tree is shown in **Fig. 4**. The most striking finding is that the placement of *Psy* isolates in the Gvg phylogeny is inconsistent with the species tree. Additionally, using the approximately unbiased (AU) and Shimodaira-Hasegawa (SH) tests with 10,000 resamplings, we determined that these two phylogenies had significantly different topologies (p < 0.01 for both tests). While we are unable to determine the origin of these gene loci in the *Pseudomonas* spp., this pattern is indicative of two or more HGT events within and/or between *gvg*-encoding *Pseudomonas* spp.

**Fig. 4.**
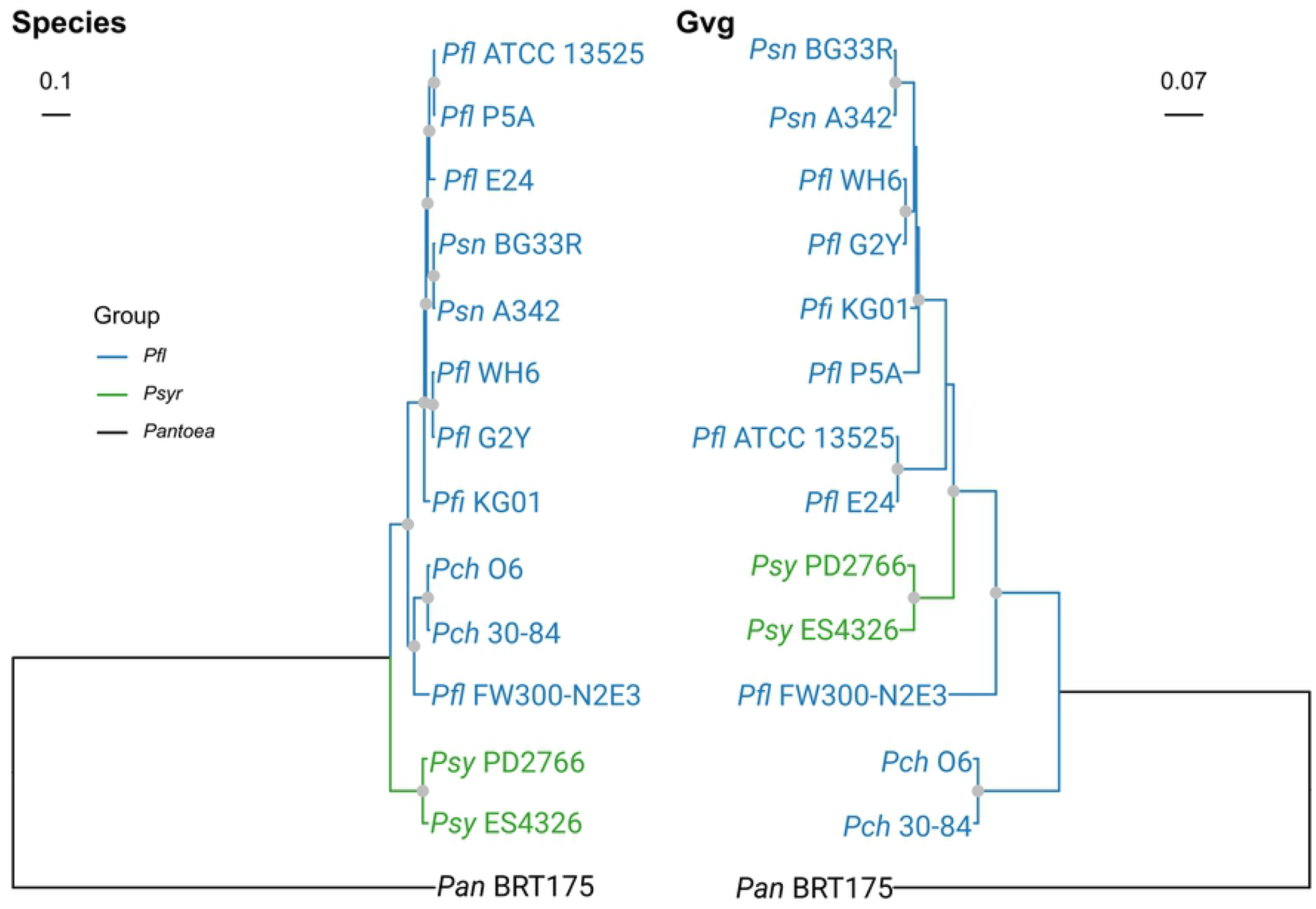
The *Pseudomonas* species and Gvg trees are incongruent. Trees were generated using maximum likelihood analysis. The species tree was inferred from the amino acid sequences of 99 housekeeping genes, while the Gvg tree was generated using a partitioned analysis of seven of the Gvg protein sequences. The trees were rooted to *Pantoea ananatis* BRT175. Nodes indicated with a gray circle are well supported (>95% UF-bootstraps & >80% SH-aLRT replicates). *Pan, Pantoea ananatis*; *Psy, Pseudomonas syringae*; *Pch, Pseudomonas chlororaphis*; *Pfl, Pseudomonas fluorescens*; *Pfi, Pseudomonas fildensis; Psn, Pseudomonas synxantha*

### Genomic context of the *gvg* locus

The *gvg* cluster appears in several distinct genetic contexts **(Fig. 3)**. These different genetic contexts correspond to both phylogeny and gene content of clusters, i.e. clusters in closely related strains have the same gene content and are located in the same genomic context **(Fig. 3)**. The genetic context is the same for all strains in the *P. syringae* group, while there are six observed in the *P. fluorescens* group (**Fig. 3)**. Strains that are closely related to *gvg*-cluster-containing strains but that do not contain *gvg* clusters maintain synteny of the neighboring genes. This suggests specific insertion of *gvg* clusters rather than larger genetic rearrangements. We propose that each example of a cluster located in a distinct genetic context represents an independent insertion of the *gvg* cluster in that lineage.

There is one example where *gvg* clusters are present in the same genetic context even though the strains are in multiple clades of the phylogeny. These strains consist of members of the *Pfl* WH6 clade, the *Psn* A342 clade and *Pfl* 1112. In all of these strains, the *gvg* cluster is located in the same chromosomal location, between *tam* and *ssb* genes. The *tam* gene encodes a methyl transferase and *ssb* encodes a single stranded DNA-binding protein. Remarkably, this common location of the *gvg* cluster is true despite the many related strains lacking the cluster **(Fig. S3)**. Two possible explanations for this observation are that 1) the gene cluster has been inserted in at least three lineages in the same location or 2) the common ancestor to these three clades contained the *gvg* cluster at this location but that it has been lost in multiple lineages. In strains without the *gvg* cluster at this location, alternative genes present between *tam* and *ssb* encode a LysR transcription factor, chloroperoxidase, chemotaxis protein, and various hypothetical proteins. In some strains, an integrase, transposase or restriction enzyme is also apparent. Like the presence of the *gvg* cluster, these variations are only partially consistent with phylogeny **(Fig. S3)**. For example, chloroperoxidase and *lysR* regulatory proteins are present in this location in several disparate clades **(Fig. S3)**. This analysis reveals that the position between the *tam* and *ssb* genes is highly variable in the *P. fluorescens* subgroups, perhaps indicating a hotspot for recombination or insertion.

These observations regarding HGT of the FVG biosynthetic cluster are consistent with the evidence of HGT for other specialized metabolites produced by plant-associated bacteria, including several metabolites commonly associated with *Pseudomonas* (reviewed in [51,52]). For example, the gene clusters responsible for biosynthesis of 2,4-diacetylphloroglucinol (DAPG), phenazines and pyochelin are observed in *Pseudomonas* as well as disparate, unrelated taxa [53–56]. As we observed with the *gvg* cluster, biosynthetic gene clusters frequently are present in both *Pseudomonas* and *Burkholderia* in particular, which may be explained by the presence of these bacteria in the same habitats [51].

Similarly, the vinylglycine rhizobitoxine is produced by bacteria from multiple genera, including the rhizobial *Bradyrhizobium elkanii* and the plant pathogen *Burkholderia andropogonsis*. Rhizobitoxine biosynthesis (*rtx*) genes flanked by insertion sequences have been reported in *Xanthomonas oryzae* pv. *oryzae* KACC10331 [57], though rhizobitoxine production has not been reported in this strain. AMB production has only been reported in *Pseudomonas aeruginosa* and AVG in one strain of *Streptomyces* sp., though systematic analyses have not been performed for other vinylglycines.

### Characteristics of *gvg*-like gene clusters

In addition to the strains containing *gvg* gene clusters, there are many strains with only some of the genes of the cluster, sometimes in combination with different genes. These variations are present in a broader range of bacteria than is the complete *gvg* cluster **(Table S4)**. Genera with strains containing these *gvg*-like clusters include the Alphaproteobacterium *Salinarimonas*; Betaproteobacteria *Burkholderia, Ralstonia* and *Thiomonas*; Gammaproteobacteria *Pseudomonas, Pantoea, Acinetobacter* and *Serratia*; and the Actinobacteria *Streptomyces* and *Saccharopolyspora* **(Table S4)**. These clusters contain genes in various arrangements and do not contain all of the genes shown to be necessary for FVG production [17], and thus would not be predicted to produce FVG. The core genes of these cluster variants are *gvgA, gvgB, gvgC*, and *gvgF*. The vast majority of these gene clusters are uncharacterized.

The evolutionary relationship of the *gvg* and *gvg*-like clusters was examined by constructing a phylogeny of the carbamoyltransferase (CTase) encoded by *gvgF*, which is common to the majority of the gene clusters, including the complete *gvg* clusters discussed above. The CTases from gene clusters with the same gene content tend to be in the same clades, regardless of taxonomy **(Fig. 5)**. Additionally, the CTases from the complete *gvg* clusters form a separate clade from those in *gvg*-like clusters, irrespective of taxonomy. (The exception is CTases encoded by truncated clusters from the *Burkholderia mimosarum* strains, which are also found in this clade, suggesting a relatively recent origin from a complete *gvg* cluster.) **(Fig. 5)**.

**Fig. 5.**
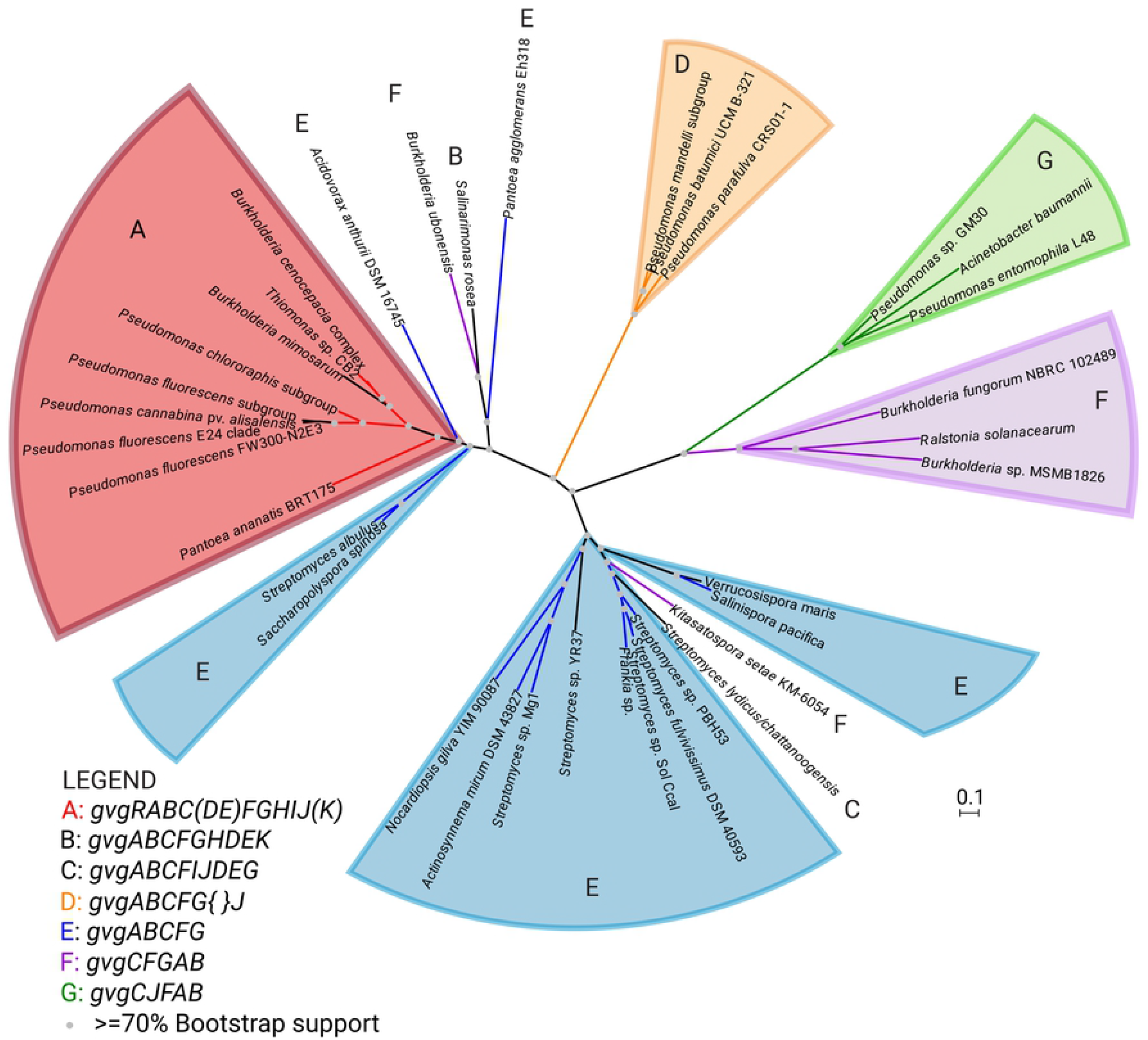
The structure of the *gvg* gene cluster is correlated with the inferred *gvg* phylogeny. This is a maximum likelihood phylogenetic tree of the carbamoyltransferase encoded by *gvg* clusters and other gene clusters. Clades are colored and labeled based on the genes present in the gene cluster. The red A clade corresponds to a complete *gvg* cluster with all of the genes present for production of FVG. In the key, the genes present in each cluster are listed, with parentheses around genes that are optional and curly braces indicating insertions. Sequence accessions are listed in **Table S5**.

At least eight cluster variations are present **(Fig. 5)**, leading to the hypothesis that each contributes to the production of a unique compound, perhaps an amino acid or similar product. One variant is located within a known cluster for biosynthesis of the macrolide antibiotics aldgamycin and chalcomycin in *Streptomyces* sp. HK2006-1 [58]. This cluster variation contains homologs of *gvgA, gvgC, gvgF* and *gvgG*, annotated as *almUII, almUIII, almDIV* and *almUIV*, respectively. However, exactly how these genes may function in the synthesis of the cluster products is unknown. These *gvg*-like loci are promising candidates for future molecular characterization. An important question is whether these gene clusters lead to the production of as yet undiscovered compounds or to previously identified compounds for which the biosynthetic genes are not yet known.

The presence of subsets of core biosynthetic genes in unrelated strains has also been observed for the phenazine *phz* and DAPG *phl* clusters encoded in other *Pseudomonas* spp. [49, 50]. This recycling of biosynthetic sub-clusters into new clusters may be a key method for evolution of biosynthetic diversity [59,60].

### CONCLUSION

Historically, secondary metabolites were identified in single organisms or groups of organisms. The range of organisms capable of producing the metabolites were not known. With the advent of genome sequencing and the linking of genes clusters to metabolites, it is now possible to obtain a deeper understanding of their distribution and abundance.

Analysis of sequence databases reveals that the *gvg* cluster is widespread within the genus *Pseudomonas*. It appears in strains commonly studied and utilized for biological control of a number of pathogens/pests, including all sequenced strains of *P. chlororaphis*. FVG was produced and excreted by both strains of *P. chlororaphis* tested and its activity may be complementary to other useful properties of these strains. Production of FVG is not a defining characteristic of the *P. fluorescens* group, as only some of the strains are capable of producing the compound. The *gvg* cluster is not restricted to single clade or subgroup of the *P. fluorescens* group, but instead is dispersed throughout the group in multiple subgroups. This varied pattern indicates that FVG provides a function that is beneficial in specific ecological contexts and niches but may be a burden in others, leading to loss of the *gvg* locus even in closely related isolates.

In contrast, although the *gvg* cluster is present in several pathogenic strains of the *P. syringae* group, FVG was not produced under the conditions tested for three of the strains. Additional cues for the induction of FVG production may be required in these strains. Experiments to understand different modes of regulation of FVG production in these strains are ongoing. However, FVG production is not restricted to *Pseudomonas*. The strain *Pan* BRT175 was confirmed to produce FVG. In this case, FVG biosynthesis must be embedded in different regulatory networks than in *Pseudomonas*. The unexpected, diverse pattern of *gvg* encoding strains indicates that FVG and related compounds may have additional benefits and outcomes that have yet to be elucidated.

The pattern of distribution of *gvg* cluster, non-congruence between gene and species trees, and variation in genetic context suggest that the cluster has been horizontally transferred multiple times. In some cases, indicators of integration consistent with a “classic” genomic island, such as an integrase, transposase or tRNA genes, are located adjacent to the cluster. However, the lack of these indicators in many strains may be reflective of an ancient integration event, for which those signs are not typically found. The frequency of which the cluster appears in unrelated strains suggest a useful function for a variety of bacteria living in a variety of plant or soil-related environments.

Finally, *gvg*-like gene clusters containing some of the genes of the *gvg* clusters are found in a wide range of bacteria. Study of these gene clusters may help us to understand the biosynthetic pathway of FVG as well as discover novel compounds or result in new sources of known compounds.

## ACKNOLWEDGEMENTS

We thank Anne Anderson, Carolee Bull, Jeff Chang, Daniel Kluepfel, Joyce Loper, Leland Pearson, John Stavrinides, and Pat Wechter for generously providing bacterial strains. Strain P5A was isolated during a classroom-based research experience by students in the class of Alice Eldridge at Philomath High School. The LAESI-MS analysis was performed by Callee Walsh and Gregory Boyce at Protea Biosciences, Inc.

## SUPPORTING INFORMATION

**Fig. S1. The *gvg* gene cluster maintains co-linearity, including outside of the *Pseudomonas* genus**. Comparison of *gvg* gene clusters in *Pseudomonas fluorescens* WH6, *Pantoea ananatis* BRT175, *Thiomonas* sp. CB2 and *Burkholderia cenocepacia*. Gene arrows are colored by function and are to scale. Gray arrows indicate genes not present in *gvg* clusters from *P. fluorescens*.

**Fig. S2. Promoter regions are different between *Pseudomonas fluorescens* and *P. syringae***. Comparison of the promoters in the intergenic region between *gvgR* and *gvgA* of the *gvg* clusters from *Pseudomonas fluorescens, P. chlororaphis* and *P. syringae* strains. The locations of the -35 and -10 promoter sites for *P. fluorescens* WH6 are indicated by gray arrows above the sequences.

**Fig. S3. The *gvg* locus inserted in a putative hotspot for horizontal gene transfer in some isolates**. This is a subset of **Fig 2** focusing on the *Pseudomonas fluorescens* subgroup. The putative identity of genes present between the *tam* and *ssb* genes in a given strain are indicated by color (red = *gvg* locus; turquoise = *lysR* protein; green = chloroperoxidase protein; orange = chemotaxis protein; purple = other genes; gray = no insert). Strains containing a *gvg* locus are shown in bold.

**Table S1. Strains used to construct multi-locus sequence phylogeny of Pseudomonas**.

**Table S2. Strains containing gvg biosynthetic gene cluster**.

**Table S3. LAESI-MS ablation values for FVG (m/z 183.037)**.

**Table S4. Bacterial strains with gvg-like clusters**.

**Table S5. Carbomoyltransferase protein sequences used to construct phylogeny.**

**File S1. Full maximum likelihood phylogenetic tree**.

